# Within-colony microbial response of three species with different susceptibility to Stony Coral Tissue Loss Disease

**DOI:** 10.64898/2026.06.09.726945

**Authors:** J. Eduardo Aguayo-Leyva, Zita P. Arriaga-Piñón, Lorenzo Álvarez-Filip, Anastazia T. Banaszak, José Q. García-Maldonado, David A. Paz-García

## Abstract

Stony coral tissue loss disease (SCTLD), a coral pathology with rapid tissue loss and high mortality rate has affected nearly 30 species with a variable degree of susceptibility across species. It has been suggested that SCTLD has a systemic effect within coral colonies, but little is known about within-colony changes of the microbial communities associated with this disease. Here we evaluate the microbial shifts within coral colonies: apparently healthy tissue and SCTLD tissue. The study was done in three species following a gradient of susceptibility to the disease: *Dendrogyra cylindrus* (Dcyl, n = 11) and *Pseudodiploria strigosa* (Pstr, n = 6) two highly susceptible species; and *Orbicella faveolata* (Ofav, n = 8), a moderately affected species. 16S rRNA Illumina sequences analysis showed differential microbial community structure within two species (Dcyl, p = 0.01, Pstr, p = 0.01) but not for Ofav (p = 0.28). Taxonomic profiles of bacterial groups were species-specific in SCTLD tissue, but healthy tissue shared similarities between species including Pirelullales, NB1-J and SAR324. Our results reveal that the microbial communities’ effects associated to the disease follow a similar pattern to the species susceptibility to SCTLD, providing new insights into the disease dynamics in the Mexican Caribbean.

## Introduction

Corals harbor multiple symbiotic relationships between the coral host, its endosymbiotic algae (Symbiodinaceae), microeukaryotes, bacteria, archaea, fungi, and viruses [1,2], comprising what is known as the “coral holobiont”. This has a critical role in the health and ecological success of corals as it provides protection and metabolic resources [3]. Particularly, coral-associated bacteria contribute to biogeochemical cycles (e.g., C, N, S), produce secondary metabolites crucial to the coral-Symbiodiniaceae relationship, play an important role in larval recruitment and settlement and provide protection against pathogens [4]. Bacterial communities associated with corals can exhibit high variability due to the complexity of the holobiont components and to the dynamics of the coral reef environment [5], responding to several factors including ocean warming, pollution, and diseases [6,7].

Stony coral tissue loss disease (SCTLD) is a plague-type pathology that affects scleractinian corals. It is characterized by acute focal/multifocal or locally extensive tissue loss lesions that lead to the exposure of the coral skeleton [8] and can cause the mortality of a colony within weeks [9]. It was first observed along the Florida Keys in 2014 [10] and spread to the Caribbean, with the first observation of SCTLD in Mexico during the summer of 2018 [11]. SCTLD affects nearly 30 species with different degrees of susceptibility. Corals of the family Meandrinidae, such as *Dendrogyra cylindrus* and *Meandrina meandrites*, have mortality rates over 90% [12] and *D. cylindrus* has been reported as locally extinct [13]. Other highly susceptible species include the brain corals, such as *Pseudodiploria strigosa* (subfamily Faviinae) with a mortality rate between 60 and 100% [14]; while most moderately afflicted species, such as those from the genus *Orbicella* (family Merulinidae), have a lower risk although with population losses ranging between 20 and 40% [15].

Although the etiological agent is yet to be determined, previous studies have reported that a primary infection by viruses targeting Symbiodiniaceae cells [16,17] followed by a secondary infection by opportunistic bacterial pathogens, due to the evidence of the effectiveness of antibiotics in halting the lesion progression [18]. Previous reports of SCTLD microbial communities indicate that the pathology could induce systemic effects due to the histological evidence of lesions in healthy tissue in affected colonies, higher alpha diversity of the microbiome prior to the development of lesions [19], and no significant differences in composition and structure between affected and healthy tissue in *Colpophyllia natans* colonies [20]. Analyses of the microbial communities using 16S rRNA have identified bacterial taxa associated with SCTLD lesions, including Rhizobiales, Clostridiales, Peptostreptococcales-Tissierellales, Rhodobacterlaes, Flavobacterales and Vibrionales [21–24]. Despite the progress in the knowledge of the impacts of SCTLD in scleractinian corals, little is known about the effects between SCTLD-affected and apparently healthy tissue within diseased colonies and its association to the degree of susceptibility of coral host species. The present study aims to characterize the within-colony effects on the structure and composition of the microbial communities of three coral species that are moderately to highly susceptible to SCTLD in the Mexican Caribbean.

## Material and methods

### Sample collection

This study was undertaken in the Mexican Caribbean and focused on three coral species: *Dendrogyra cylindrus* (Dcyl) and *Pseudodiploria strigosa* (Pstr) with a high prevalence of SCTLD (80 to 100%) and *Orbicella faveolata* (Ofav) with moderate susceptibility to SCTLD (40-20%) (Alvarez-Filip et al. 2022; Mendoza Quiroz et al. 2023). Samples of *D. cylindrus* were collected from Colombia Somero, Cozumel Island in April 2019, while samples of *P. strigosa* and *O. faveolata* were collected in Tanchacté reef in the region of Puerto Morelos in August 2019. The two sampling sites are separated by an approximate distance of 68 km.

For the three species, samples were taken both from the apparently healthy area of the colony (hereafter healthy tissues) and from lesions affected by SCTLD near the edge of the colony (hereafter SCTLD tissue,). *D. cylindrus* samples were collected from 4 colonies (7 SCTLD tissues and 4 healthy tissues), *P. strigosa* samples were collected from 3 colonies (3 SCTLD tissues and 3 healthy tissues), and *O. faveolata* samples were collected from 4 colonies (4 SCTLD tissues and 4 healthy tissues). Collection was performed using a hammer using fresh gloves per sample and immediately stored in individual plastic bags. Healthy tissue samples were collected before SCTLD tissue samples to avoid cross contamination. Samples were stored on ice and were transported to the laboratory where they were transferred to 1.5 mL microtubes containing RNALater ® and stored at -20°C until further processing.

Collections were performed using fresh gloves per sample to prevent cross-contamination and healthy tissue samples were collected before the SCTLD tissue samples. Samples were placed in individual bags and stored in RNALater® and were transported to the laboratory for further processing.

### DNA isolation, amplicon library preparation and sequencing

DNA isolation, library preparation and sequencing methods were described in a previous publication [25]. Briefly, 0.2 g of coral tissue per sample was used for DNA isolation using the Qiagen DNeasy ® Blood & Tissue Kit following the manufacturer′s protocol. DNA integrity was verified by agarose gel 1% electrophoresis. PCR amplification of V3 and V4 regions of the 16S rRNA gene was carried out according to the Illumina metagenomic sequencing library preparation, using 341F and 805R primers [26]. A no-template control was included in each PCR run for monitoring contamination. Indexed PCR products were purified using AMPure XP (Beckman Coulter ®) magnetic pearls and were quantified using Qubit ® Fluorometer to verify the size of the library. Sequencing was carried in the Illumina-MiSeq platform with the MiSeq reagent kit V2 (2×250 pb) following the manufacturer’s instructions. Sequencing was performed in the Aquatic Pathology lab at Cinvestav Mérida.

### Analysis of amplicon libraries

Paired and demultiplexed sequences were processed in the QIIME 2 platform (version 2024.2) [27]. After manual inspection, forward and reverse reads were trimmed at position 40 from the 5’ end and truncated at position 240 from the 3’ end to filter out reads with low quality. Denoising, error correction, and chimera removal was performed using the DADA2 plug-in [28] to finally resolve the Amplicon Sequence Variants (ASVs). Taxonomic assignment of representative ASVs was done using the V-SEARCH plug-in using the SILVA version 138 [29] database as a reference. Representative sequences were aligned using the MAFF method [30] and a phylogenetic tree was constructed using FastTree 2 [31]. The ASV table and the phylogenetic tree were exported to the R environment (version 4.1.1) [32]. The ASV table was filtered to delete those ASVs classified as mitochondria, chloroplast and those with an unassigned domain. An abundance table was then normalized using a cumulative sum scaling (CSS) transformation using *phyloseq_transform_css* from the metagMisc library [33]. Statistical analysis and visualization of results were perfomed using *phyloseq* version 1.48.0 [34], *vegan* version 2.6-6.1 [35] and ggplot2 [36] libraries.

Differences in microbial communities were assessed between healthy and SCTLD tissues. The Shannon diversity index was estimated [37]. A principal coordinate analysis (PCoA) was performed using the Bray-Curtis distance to generate the ordination plot to assess structural differences in the microbial communities. Permutational analysis of variance (PERMANOVA) was carried out to detect structural differences in the microbial communities. To detect differences in the taxonomic composition of microbial communities, linear discriminant analyses with Effect Size (LefSe) were performed at the order level using a cutoff of LDA > 2 and a p-value <0.05 for the internal Kruskal-Wallis and Wilcoxon test using the *microeco* package (ver. 1.8) [38].

## Results

### Within-colony differences in microbial communities between healthy and SCTLD tissues

The analysis of the microbial communities showed distinct within-colony structure between healthy and SCLTD tissues in the three studied coral species (figure 1). The ordination (PCoA) of amplicon sequence variants showed distinct compositional profiles and consistent separation between healthy and SCTLD tissues within a single colony (figure 1b). Significant differences between tissues were supported in *D. cylindrus* (F_1,7_ = 1.56, p = 0.01) and *P. strigosa* (F_1,2_ = 2.33 p = 0.01), but not for *O. faveolata* (F_1,5_ = 1.04, p = 0.2877). These results suggest that coral tissue lesions are associated with significant alterations in the coral microbiome composition at the within-colony level, regardless of the SCTLD prevalence of the host species.

**Figure 1a.**
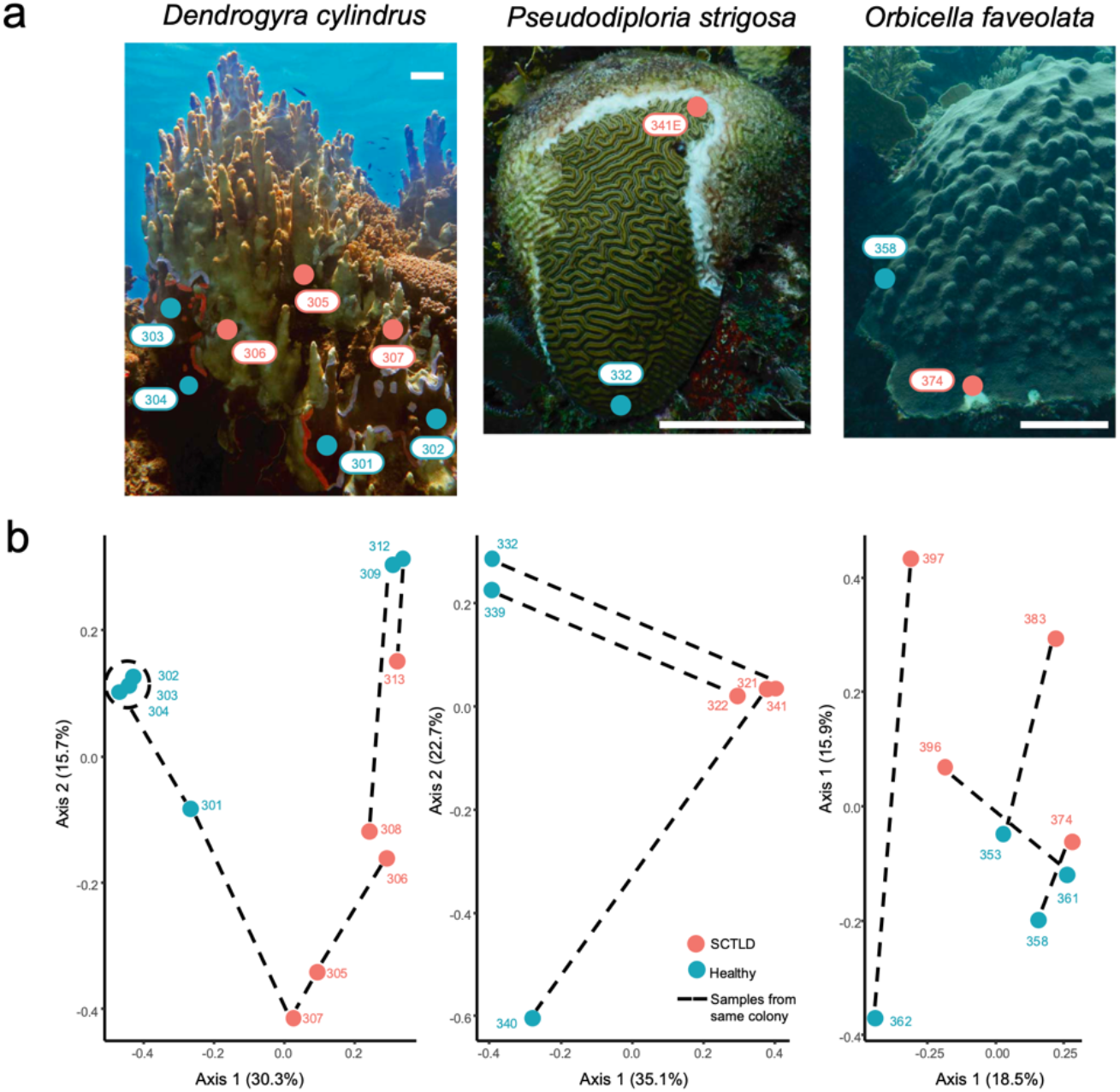
Example of colonies of *Dendrogyra cylindrus, Pseudodiploria strigosa* and *Orbicella faveolata*. Scale bar equals 20 cm. **1b** Principal coordinate analysis (PCoA) ordination plots of amplicon sequence variants from apparently healthy (blue) and SCTLD-affected samples (red). Dashed lines connect samples from the same colony.

### Taxonomic profiles and differentially abundant taxa explain within-colony differences in microbial communities

All the samples of *Dendrogyra cylindrus* were predominantly dominated by Oceanospirillales, Rhizobiales, Vibrionales, and SAR86 (figure 2a). Oceanospirillales was a predominant group that showed an increase in relative abundance from 10.61% in healthy to 34.31% in SCTLD tissues. Additionally, several low-abundance orders (< 2% relative abundance) showed varying patterns of enrichment or depletion in SCTLD tissue, contributing to the overall compositional shifts observed in the diseased state. The orders in *D. cylindrus* samples that were detected as differentially abundant (figure 2b) for healthy tissue included Pirelullales, Verrucomicrobiales, Steroidobacterales, Nostocales, Rickettsiales, Opitulales and the SAR324 clade (Marine group B), while in the SCTLD tissue samples were represented by the Rhizobiales and Micrococcales orders.

**Figure 2a.**
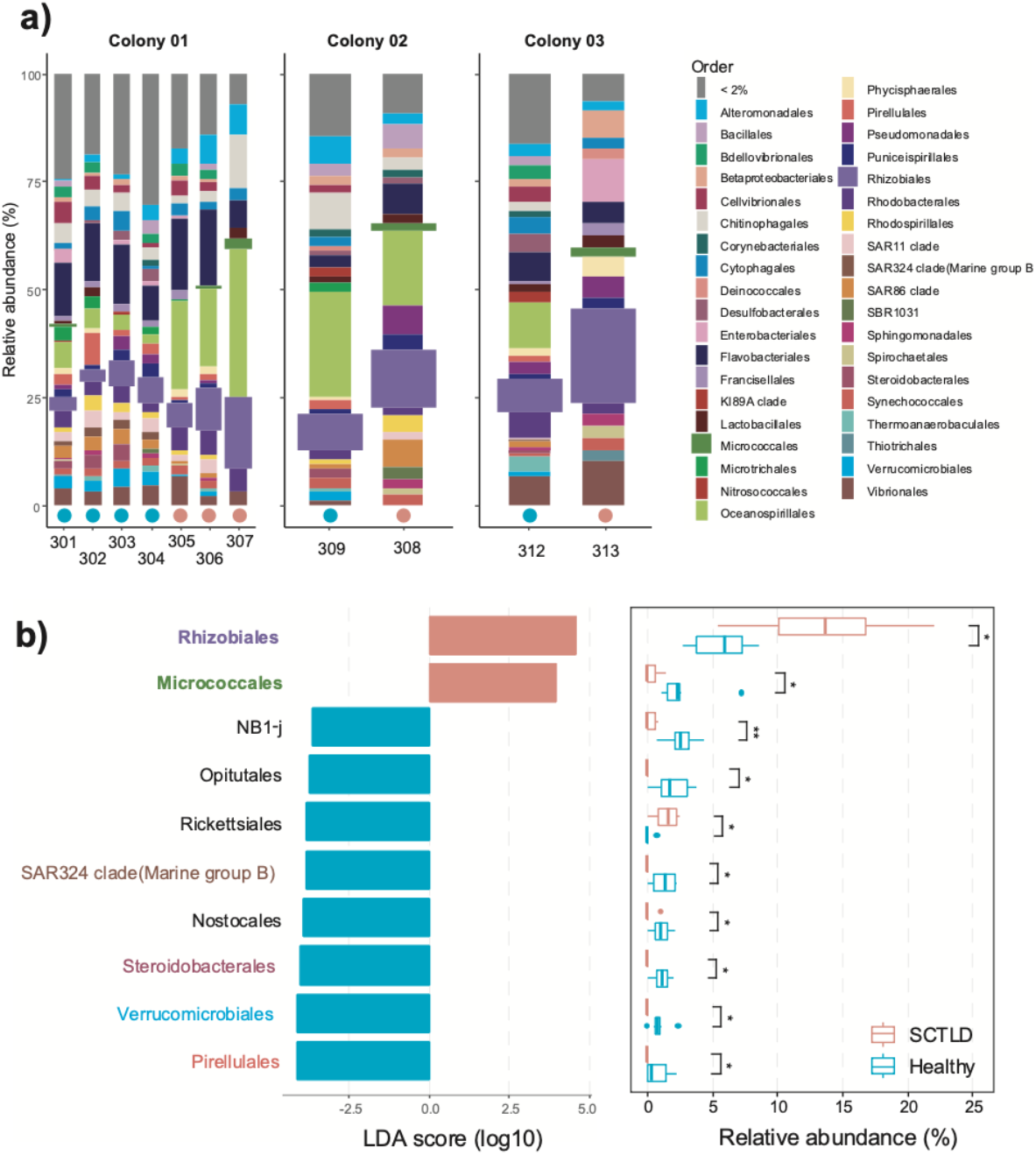
Microbial community composition of samples from three colonies of *Dendrogyra cylindrus*. Healthy tissue is indicated with a blue circle and SCTLD tissue with a red circle. Differential SCTLD abundant orders are shown as wider columns. **2b** Linear discriminant analysis with an effect size (LEfSe) plot with differential abundant orders by condition and their LDA scores. Significance level: *: P <0.05; **: P <0.01

For *Pseudodiploria strigosa*, the taxonomic profiles demonstrated that only a few groups represented more than 75% of relative abundance in SCTLD than in healthy tissue (figure 3a). Ten important orders with high relative abundance and with differential abundance in the SCTLD tissues were detected: Vibrionales (0.43 to 29.92%), Peptostreptococcales-Tissierellales (22.42%), and Desulfovibrionales (6.44%). Peptostreptococcales-Tsisserellales, Campylobacterales, Desulfovibrionales, Bacteroidales, Clostridiales and Oceanospirillales (figure 3b). By contrast, fourteen orders were found to be differentially abundant in healthy samples: Rhizobiales, Pirellulales, Cytophagales, Rhodobacterales, Kiloniellales, Microtrichales, Steroidobacterales, the clade NB1-j, Polyangiales, Verrucomicrobiales, Thermoanaerobaculales, Bdellovibrionales, Thalassobaculales and Caulobacterales (figure 3b).

**Figure 3a.**
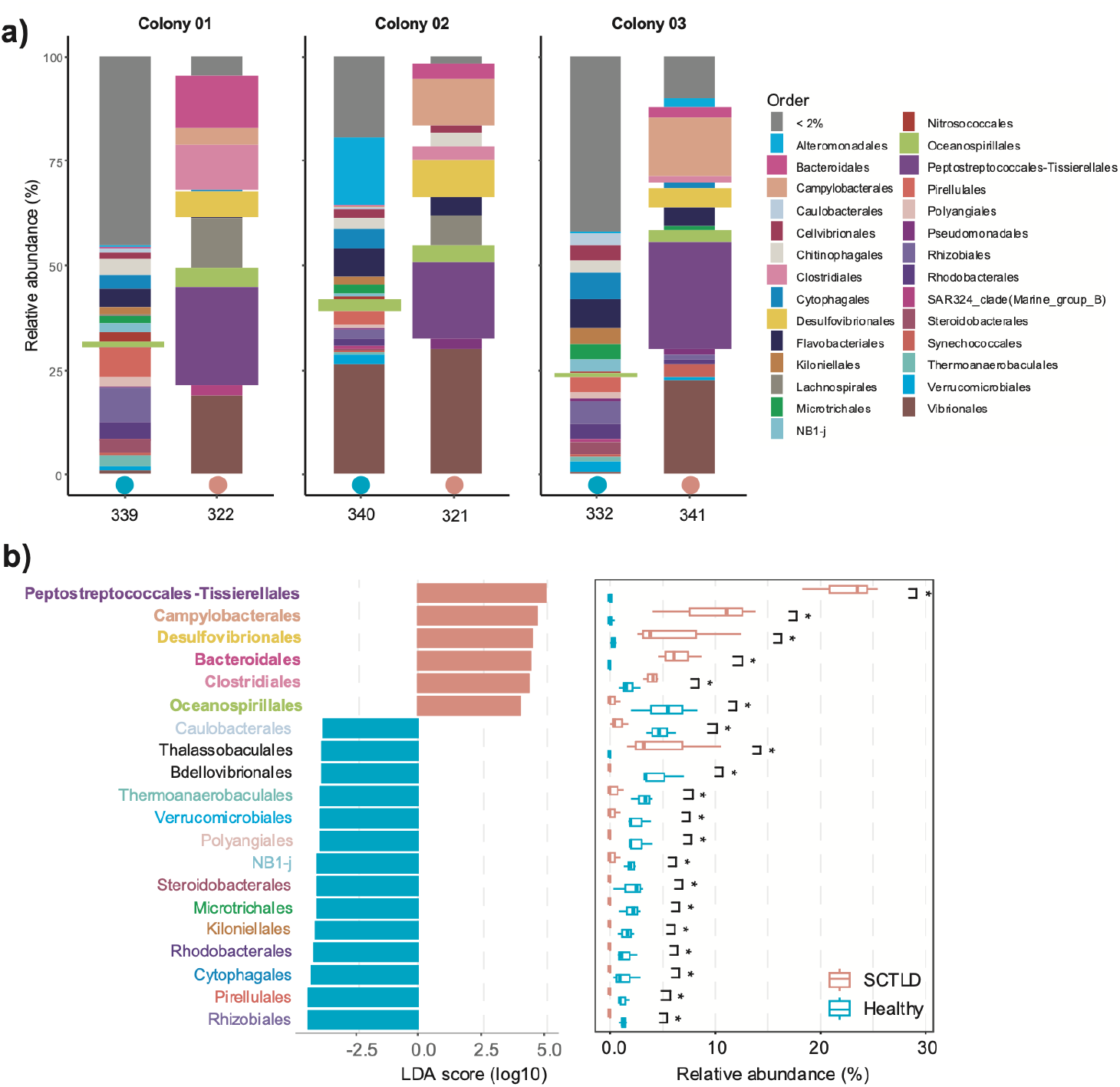
Microbial community composition of samples from three colonies of *Pseudodiploria strigosa* at the order level. Healthy tissue is indicated with a blue circle and SCTLD tissue with a red circle. Differential SCTLD abundant orders are shown as wider columns. **3b** Linear discriminant analysis with effect size (LEfSe) plot with differential abundant orders for by condition and their LDA scores. Significance level: *: P <0.05

For the *Orbicella faveolata* samples, the taxonomic profiles showed that the most well-represented groups in healthy tissue were Chitinophagales (6.25%), Cythopaghales (5.8%) and Flavobacteriales (5.38%). By contrast, in the SCTLD tissue, Flavobacteriales (8.7%), Cytophagales (6.22%), and Alteromonadales (6.92%) were the most abundant taxa (figure 4a). Differential abundance analysis for the species (figure 4b) indicated that Cyanobacteriales and Haliangiales were the taxa that characterized the SCTLD tissue samples, while Planctomycetales, the SAR324 clade (Marine group B) along with the clades Subgroup-9 and 0319-6G20 were the most representative taxa for healthy tissue.

**Figure 4a.**
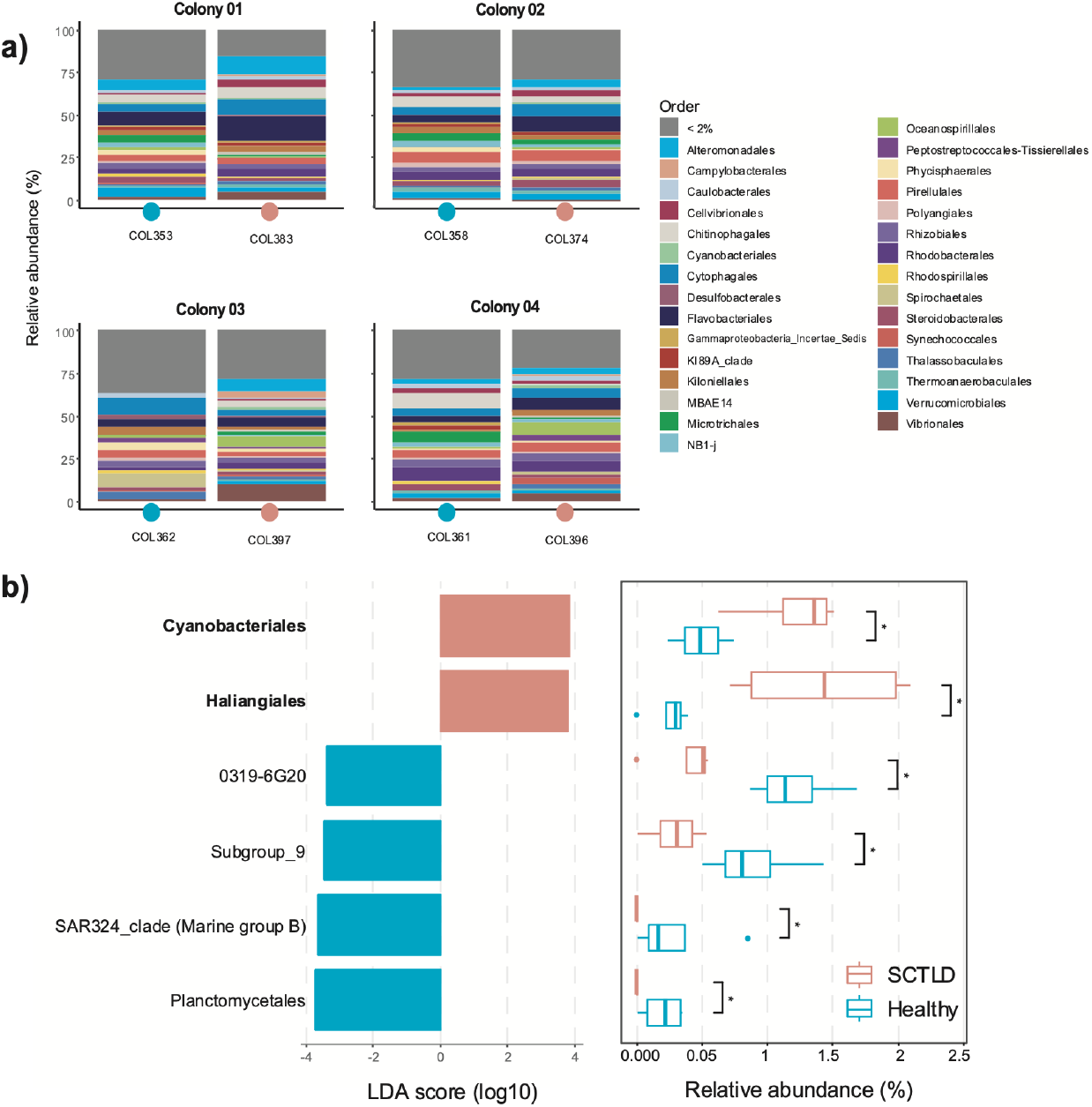
Microbial community composition of samples from four colonies of *Orbicella faveolata* at the order level. Healthy tissue is indicated with a blue circle and SCTLD tissue with a red circle. Differential SCTLD abundant orders are shown as wider columns. **4b** Linear discriminant analysis with effect size (LEfSe) plot with differential abundant orders for by condition and their LDA scores. Significance level: *: P <0.05

## Discussion

The present study shows evidence of within-colony response on the microbiome of three species with different degrees of susceptibility to SCTLD in the Mexican Caribbean. Within-colony assessment of healthy and SCTLD tissues revealed species specific microbial responses, structural and compositional differences in highly susceptible coral species (*D. cylindrus* and *P. strigosa*), whereas the moderately susceptible species (*O. faveolata*) showed no significant differences in its microbial communities. These findings indicate that coral species susceptible to SCTLD demonstrate distinct microbiome responses at the within-colony level, suggesting a localized effect of the disease. Differential exposure of corals to environmental factors such as light intensity, water flow, nutrient gradient and exposure to disease can vary within a single coral colony [39]. These differences may create microhabitats that can modify microbial community dynamics, increasing the complexity of the coral microbiome. Thus, the microbiome can be differentially responsive depending on the position within a single colony (i.e. within-colony differentiation) to mitigate the effects of stress such as high temperature [40], sedimentation [41], and diseases [42].

### Within-colony microbiome difference depends on species susceptibility to SCTLD

Coral species of the families Meandrinidae and Faviidae are the most highly susceptible and affected corals by SCTLD, while members of the Merulinadae are of moderate susceptibility [15,43]. Differential lesion progression rates have also been observed, as *P. strigosa* has exhibited ranges of tissue loss up to 25 cm^2^/day while for *Orbicella* species this rate was estimated to be around 3 cm^2^/day [44]. Although progression rate may vary geographically, species consistently exhibit similar patterns of lesion progression rates and mortality by SCTLD. The microbiome of *D. cylindrus* (Meandrinidae) and *P. strigosa* (Faviidae) showed within-colony differentiation related to the tissue condition (healthy tissue vs. SCTLD tissue, figure 1), while the response of *O. faveolata* (Merulinidae) was less evident.

Previous studies have reported microbial shifts between healthy and SCTLD affected colonies of *P. strigosa* [45] and *O. faveolata* [19–21], while this study is the first that describe the microbiome of *D. cylindrus*. Overall, reports on the microbial communities in both affected and unaffected tissues suggest a systematic widespread effect of the disease on the microbiome within each colony. In contrast, our results suggest a localized response and differentiation of the microbial communities (i.e., within-colony response) dependent on the species susceptibility to SCTLD. Additionally, the results of this study indicate a within-colony effect related to tissue health. For example, healthy tissues of the *D. cylindrus* colony (samples 302—304) remained closely clustered in comparison to the SCTLD tissue (305—307, Fig. 1b) exhibiting similarities in the microbial communities’ structure.

Within-colony differences between SCTLD and healthy tissue were related to the degree of susceptibility to the disease of the coral species. There are hypotheses about how the differences in coral species susceptibility to SCTLD can, in turn, modify the microbiome. First, previous reports have stated that SCTLD is a disease that is transmissible by direct contact of tissue, but water flow and sediments can also act as vectors [24,46]. Given that bigger colonies with horizontal or planar surfaces tend to be more exposed to water and retain more sediment than other colony morphologies [47], *D. cylindrus* and *O. faveolata* species would be expected to be more susceptible to SCTLD. Second, Symbiodiniaceae species harbored by the coral host exhibit a gradient of vulnerability, with *Breviolum* being the most affected, followed by *Cladocopium, Durusdinium* and *Symbiodinium* [48]. It has been reported that *D. cylindrus* has high symbiont specificity with *Breviolum* being the dominant group (up to 80% abundance) associated with this coral species [49]. Endosymbiotic profiles of *P. strigosa* showed dominance and stability with *Breviolum* followed by *Cladocopium*, in comparison to *O. faveolata*, which has an exclusive association with the *Cladocopium* genus [50]. The disruption of Symbiodiniaceae cells could result in alterations of the bacterial component of the holobiont as both elements of the holobiont interact within the colony [17].

### Within-colony effects of SCTLD in microbial taxa are species specific

Our results showed that there were some similarities between microbial profiles in SCTLD tissue across species, including bacterial taxa that has been previously identified as enriched in diseased tissue, such as Rhizobiales, Clostridiales, Bacteroidales, Oceanospirillales and Vibrionales [19,20]. These similarities may be related to those groups that are known to be opportunistic bacteria that can necrotize host tissue [51]. As diseased states can alter the microbiome [2], affected corals can harbor opportunistic pathogens, secondary colonizers and saprophytic bacteria, resulting in increasing microbiome diversity [52]. Thus, the microbiome composition tends to be species specific [53], but diseases can alter that specificity. SCTLD tissue of *D. cylindrus* (figure 2a) had the presence of Micrococcales from the phylum Actinobacteria, whose role in coral health has not been documented, but it has been suggested that it could produce secondary metabolites to control microbial communities in mammals [54]. Flavobacteriales and Cytophagales are also abundant bacterial groups in SCTLD tissue and have been previously reported as enriched taxa in diseased tissue [20] and are known for the potential production of biofilm layers that could lead to the colonization of other bacteria [55].

In SCTLD tissues of *P. strigosa* there was an abundance of Peptostreptococcales-Tissierellales (figure 3a). Bacteria from this order have been previously reported as abundant in bleached tissue in corals and enriched in SCTLD tissue. Their potential role in the disease has been linked to their production of toxins that degrade Symbiodiniaceae cells (Becker et al. 2021).

Differential analysis of ASVs for *O. faveolata* (figure 4b) revealed that the groups characterizing both health conditions were in low relative abundance. Cyanobacteriales was associated with SCTLD-tissue, this order has previously been linked to coral tissue lysis [56], but also with the production of antimicrobial compounds used as a treatment for Black Band Disease [57]. Haliangiales is a poorly studied bacterial order but has been associated with grey patch disease in Micronesia [58].

Healthy tissue of *D. cylindrus* (figure 2a) and *P. strigosa* (figure 3a) shared some similarities, with Pirelullales being a well-represented order for both species. This bacterial group has been correlated with wastewater and anaerobic ammonium oxidation [59]. SAR324, previously known as Marine Group B is a diverse marine order and metabolically versatile and not previously correlated with corals nor with diseases [60], but it was represented in both *D. cylindrus* and *O. faveolata*. Another group that was well-represented, specifically in *P. strigosa* healthy samples was Rhodobacterales associated with diseased states in coral reefs [61]. This order is of particular interest as it was described as one of the key biomarkers for SCTLD-affected corals [22], and the presence of this group might indicate that the microbiome was showing early signs of a diseased state.

Some of the groups present in SCTLD tissue are also present in healthy samples across the studied species. This could be explained through the following hypotheses. First, some of the detected bacterial orders are significatively diverse and can harbor multiple metabolisms and functions. Second, some of these bacterial taxa can be part of the pathobiome, bacteria that are present on the holobiont but only become pathogenic under specific environmental conditions and host susceptibility [52]. Lastly, some bacterial taxa can exhibit plastic functionality mechanisms such as horizontal gene transmission [62].

The present study faced challenges regarding the experimental design and sample size. Studies of wild diseases are bound by the epidemiology and dynamics of the disease and its transmission, making the selection of the individuals rather complicated [63,64], this effect can be reflected in biased conclusions as they might not be representative of the population.

At the colony level, corals have displayed within-colony heterogeneity in holobiont diversity. Evidence of this pattern has been reported when studying partially bleached colonies, which may be related to within-colony Symbiodiniaceae diversity as some of the endosymbiont species are more susceptible to heat stress and irradiance exposure [65]. Also, it has been demonstrated that microenvironments can be formed due to the differential exposure of the coral polyps to a gradient of diverse physicochemical factors like water flow, temperature and nutrients and oxygen concentrations [66,67]. On the other hand, sample size is an important factor to microbiome studies as a larger dataset is required to minimize the levels of type I and II errors [68,69].

## Conclusion

The present study identified changes in the microbial communities associated with three species affected by SCTLD in the Mexican Caribbean. At the within-colony level, microbial communities exhibited differences between tissue health condition (SCTLD vs healthy tissue) and not related to the colony of origin, suggesting a localized effect of the disease rather than a systemic-level effect as in previous studies. Microbial shifts followed a gradient of susceptibility, being more disturbed in *D. cylindrus*, followed by *P. strigosa* and *O. faveolata*. Bacterial taxonomic profiles showed a degree of similarity in SCTLD tissues between samples, but changes in dominances of certain bacterial groups were detected among studied coral species. Future coral microbiome studies should consider within-colon differences as an important factor of diversity as it might reveal patterns that have not been described. These results contribute to the knowledge of the bacterial communities of corals affected by SCTLD in the Mexican Caribbean, providing information on the microbiome of *D. cylindrus*, a virtually locally extinct species.

## References

1. Hernandez-Agreda A, Gates RD, Ainsworth TD. 2017 Defining the core microbiome in corals’ microbial soup. Trends Microbiol. 25, 125–140. (doi:10.1016/j.tim.2016.11.003)

2. van Oppen MJH, Blackall LL. 2019 Coral microbiome dynamics, functions and design in a changing world. Nat. Rev. Microbiol. 17, 557–567. (doi:10.1038/s41579-019-0223-4)

3. Peixoto RS, Rosado PM, Leite DC de A, Rosado AS, Bourne DG. 2017 Beneficial Microorganisms for Corals (BMC): Proposed mechanisms for coral health and resilience. Front. Microbiol. 8. (doi:10.3389/fmicb.2017.00341)

4. McDevitt-Irwin JM, Baum JK, Garren M, Vega Thurber RL. 2017 Responses of coral-associated bacterial communities to local and global stressors. Front. Mar. Sci. 4. (doi:10.3389/fmars.2017.00262)

5. Dubé CE, Ziegler M, Mercière A, Boissin E, Planes S, Bourmaud CA-F, Voolstra CR. 2021 Naturally occurring fire coral clones demonstrate a genetic and environmental basis of microbiome composition. Nat. Commun. 12, 6402. (doi:10.1038/s41467-021-26543-x)

6. Vega Thurber R et al. 2020 Deciphering coral disease dynamics: Integrating host, microbiome, and the changing environment. Front. Ecol. Evol. 8. (doi:10.3389/fevo.2020.575927)

7. Mohamed AR, Ochsenkühn MA, Kazlak AM, Moustafa A, Amin SA. 2023 The coral microbiome: towards an understanding of the molecular mechanisms of coral–microbiota interactions. FEMS Microbiol. Rev. 47. (doi:10.1093/femsre/fuad005)

8. Hawthorn AC, Dennis M, Kiryu Y, Landsberg J, Peters E, Work TM. 2024 Stony coral tissue loss disease (SCTLD) case definition for wildlife. (doi:10.3133/tm19I1)

9. Estrada-Saldívar N, Molina-Hernández A, Pérez-Cervantes E, Medellín-Maldonado F, González-Barrios FJ, Alvarez-Filip L. 2020 Reef-scale impacts of the Stony Coral Tissue Loss Disease Outbreak. Coral Reefs 39, 861–866. (doi:10.1007/s00338-020-01949-z)

10. Precht WF, Gintert BE, Robbart ML, Fura R, van Woesik R. 2016 Unprecedented disease-related coral mortality in southeastern Florida. Sci. Rep. 6, 31374. (doi:10.1038/srep31374)

11. Alvarez-Filip L, Estrada-Saldívar N, Pérez-Cervantes E, Molina-Hernández A, González-Barrios FJ. 2019 A rapid spread of the Stony Coral Tissue Loss Disease outbreak in the Mexican Caribbean. PeerJ 2019. (doi:10.7717/peerj.8069)

12. Alvarez-Filip L, González-Barrios FJ, Pérez-Cervantes E, Molina-Hernández A, Estrada-Saldívar N. 2022 Stony Coral Tissue Loss Disease decimated Caribbean coral populations and reshaped reef functionality. Commun. Biol. 5, 440. (doi:10.1038/s42003-022-03398-6)

13. Cavada-Blanco F, Croquer A, Vermeji M, Goergen L, Rodríguez R. 2022 Dendrogyra cylindrus. IUCN Red List of Threatened Species. (doi:10.2305/IUCN.UK.2022-2.RLTS.T133124A129721366.en)

14. Mendoza Quiroz S, Tecalco Renteria R, Ramírez Tapia GG, Miller MW, Grosso-Becerra MV, Banaszak AT. 2023 Coral affected by Stony Coral Tissue Loss Disease can produce viable offspring. PeerJ 11, e15519. (doi:10.7717/peerj.15519)

15. Papke E et al. 2024 Stony Coral Tissue Loss Disease: A review of emergence, impacts, etiology, diagnostics, and intervention. Front. Mar. Sci. 10. (doi:10.3389/fmars.2023.1321271)

16. Work TM, Weatherby TM, Landsberg JH, Kiryu Y, Cook SM, Peters EC. 2021 Viral-like particles are associated with endosymbiont pathology in Florida corals affected by Stony Coral Tissue Loss Disease. Front. Mar. Sci. 8. (doi:10.3389/fmars.2021.750658)

17. Nandi S, Stephens TG, Walsh K, Garcia R, Villalpando MF, Sellares-Blasco RI, Zubillaga AL, Croquer A, Bhattacharya D. 2025 Shifts in the microbiome and virome are associated with Stony Coral Tissue Loss Disease (SCTLD). ISME Communications (doi:10.1093/ismeco/ycaf226)

18. Neely KL, Whitehead RF, Dobler MA. 2024 The effects of disease lesions and amoxicillin treatment on the physiology of SCTLD-affected corals. Front. Mar. Sci. 11. (doi:10.3389/fmars.2024.1460163)

19. Rosales SM et al. 2023 A meta-analysis of the Stony Coral Tissue Loss Disease microbiome finds key bacteria in unaffected and lesion tissue in diseased colonies. ISME Communications 3. (doi:10.1038/s43705-023-00220-0)

20. Clark AS, Williams SD, Maxwell K, Rosales SM, Huebner LK, Landsberg JH, Hunt JH, Muller EM. 2021 Characterization of the microbiome of corals with Stony Coral Tissue Loss Disease along Florida’s coral reef. Microorganisms 9. (doi:10.3390/microorganisms9112181)

21. Meyer JL, Castellanos-Gell J, Aeby GS, Häse CC, Ushijima B, Paul VJ. 2019 Microbial community shifts associated with the ongoing Stony Coral Tissue Loss Disease outbreak on the Florida Reef Tract. Front. Microbiol. 10. (doi:10.3389/fmicb.2019.02244)

22. Rosales SM, Clark AS, Huebner LK, Ruzicka RR, Muller EM. 2020 Rhodobacterales and Rhizobiales are associated with Stony Coral Tissue Loss Disease and its suspected sources of transmission. Front. Microbiol. 11. (doi:10.3389/fmicb.2020.00681)

23. Becker C, Brandt M, Miller C, Apprill M. 2021 Stony Coral Tissue Loss Disease biomarker bacteria identified in corals and overlying waters using a rapid field-based sequencing approach. Microbiology. 24, 1166–1182. (doi:10.1101/2021.02.17.431614)

24. Studivan MS, Rossin AM, Rubin E, Soderberg N, Holstein DM, Enochs IC. 2022 Reef sediments can act as a Stony Coral Tissue Loss Disease vector. Front. Mar. Sci. 8. (doi:10.3389/fmars.2021.815698)

25. Arriaga-Piñón ZP, Aguayo-Leyva JE, Álvarez-Filip L, Banaszak AT, Aguirre-Macedo ML, Paz-García DA, García-Maldonado JQ. 2024 Microbiomes of three coral species in the Mexican Caribbean and their shifts associated with the Stony Coral Tissue Loss Disease. PLoS One 19, e0304925. (doi:10.1371/journal.pone.0304925)

26. Klindworth A, Pruesse E, Schweer T, Peplies J, Quast C, Horn M, Glöckner FO. 2013 Evaluation of general 16S ribosomal RNA gene PCR primers for classical and next-generation sequencing-based diversity studies. Nucleic Acids Res. 41. (doi:10.1093/nar/gks808)

27. Bolyen E et al. 2019 Reproducible, interactive, scalable and extensible microbiome data science using QIIME 2. Nat. Biotechnol. 37, 852–857. (doi:10.1038/s41587-019-0209-9)

28. Callahan BJ, McMurdie PJ, Rosen MJ, Han AW, Johnson AJA, Holmes SP. 2016 DADA2: High-resolution sample inference from Illumina amplicon data. Nat. Methods 13, 581–583. (doi:10.1038/nmeth.3869)

29. Quast C, Pruesse E, Yilmaz P, Gerken J, Schweer T, Yarza P, Peplies J, Glöckner FO. 2013 The SILVA ribosomal RNA gene database project: Improved data processing and web-based tools. Nucleic Acids Res. 41. (doi:10.1093/nar/gks1219)

30. Katoh K, Standley DM. 2013 MAFFT Multiple Sequence Alignment Software Version 7: Improvements in performance and usability. Mol. Biol. Evol. 30, 772–780. (doi:10.1093/molbev/mst010)

31. Price MN, Dehal PS, Arkin AP. 2010 FastTree 2 – Approximately Maximum-Likelihood trees for large alignments. PLoS One 5, e9490. (doi:10.1371/journal.pone.0009490)

32. R Core Team. 2024 R: A Language and Environment for Statistical Computing.

33. Mikryukov V, Mahé F. 2021 metagMisc: Miscellaneous functions for metagenomic analysis. (https://github.com/vmikk/metagMisc)

34. McMurdie PJ, Holmes S. 2013 Phyloseq: An R package for reproducible interactive analysis and graphics of microbiome census data. PLoS One 8. (doi:10.1371/journal.pone.0061217)

35. Oksanen J et al. 2024 vegan: Community Ecology Package.

36. Wickham H. 2016 ggplot2: Elegant graphics for Data Analysis. New York: Springer-Verlag.

37. Shannon CE. 1948 A Mathematical Theory of Communication. Bell System Technical Journal 27, 379–423. (doi:10.1002/j.1538-7305.1948.tb01338.x)

38. Liu C, Cui Y, Li X, Yao M. 2021 microeco: an R package for data mining in microbial community ecology. FEMS Microbiol. Ecol. 97. (doi:10.1093/femsec/fiaa255)

39. Eaton KR, Landsberg JH, Kiryu Y, Peters EC, Muller EM. 2021 Measuring Stony Coral Tissue Loss Disease induction and lesion progression within two intermediately susceptible species, Montastraea cavernosa and Orbicella faveolata. Front. Mar. Sci. 8. (doi:10.3389/fmars.2021.717265)

40. Kemp DW, Thornhill DJ, Rotjan RD, Iglesias-Prieto R, Fitt WK, Schmidt GW. 2015 Spatially distinct and regionally endemic Symbiodinium assemblages in the threatened Caribbean reef-building coral Orbicella faveolata. Coral Reefs 34, 535–547. (doi:10.1007/s00338-015-1277-z)

41. Duckworth A, Giofre N, Jones R. 2017 Coral morphology and sedimentation. Mar. Pollut. Bull. 125, 289–300. (doi:10.1016/j.marpolbul.2017.08.036)

42. Ward JR. 2007 Within-colony variation in inducibility of coral disease resistance. J. Exp. Mar. Biol. Ecol. 352, 371–377. (doi:10.1016/j.jembe.2007.08.014)

43. Estrada-Saldívar N, Quiroga-García BA, Pérez-Cervantes E, Rivera-Garibay OO, Alvarez-Filip L. 2021 Effects of the Stony Coral Tissue Loss Disease outbreak on coral communities and the benthic composition of Cozumel reefs. Front. Mar. Sci. 8. (doi:10.3389/fmars.2021.632777)

44. Meiling S, Muller EM, Smith TB, Brandt ME. 2020 3D Photogrammetry reveals dynamics of Stony Coral Tissue Loss Disease (SCTLD) lesion progression across a thermal stress event. Front. Mar. Sci. 7. (doi:10.3389/fmars.2020.597643)

45. Thome PE, Rivera-Ortega J, Rodríguez-Villalobos JC, Cerqueda-García D, Guzmán-Urieta EO, García-Maldonado JQ, Carabantes N, Jordán-Dahlgren E. 2021 Local dynamics of a white syndrome outbreak and changes in the microbial community associated with colonies of the scleractinian brain coral Pseudodiploria strigosa. PeerJ 9. (doi:10.7717/peerj.10695)

46. Studivan MS et al. 2022 Transmission of Stony Coral Tissue Loss Disease (SCTLD) in simulated ballast water confirms the potential for ship-born spread. Sci. Rep. 12, 19248. (doi:10.1038/s41598-022-21868-z)

47. Camacho-Vite C, Estrada-Saldívar N, Pérez-Cervantes E, Alvarez-Filip L. 2022 Differences in the progression rate of SCTLD in Pseudodiploria strigosa are related to colony size and morphology. Front. Mar. Sci. 9. (doi:10.3389/fmars.2022.790818)

48. Palacio-Castro AM, Soderberg N, Zagon Z, Cooke K, Studivan MS, Gill T, Kelble C, Christian T, Enochs IC. 2025 Elevated temperature decreases Stony Coral Tissue Loss Disease transmission, with little effect of nutrients. Sci. Rep. 15, 22261. (doi:10.1038/s41598-025-06322-0)

49. Lewis C, Neely K, Rodriguez-Lanetty M. 2019 Recurring episodes of thermal stress shift the balance from a dominant host-specialist to a background host-generalist zooxanthella in the threatened pillar coral, Dendrogyra cylindrus. Front. Mar. Sci. 6. (doi:10.3389/fmars.2019.00005)

50. Cunning R, Lenz EA, Edmunds PJ. 2024 Measuring multi-year changes in the Symbiodiniaceae algae in Caribbean corals on coral-depleted reefs. PeerJ 12, e17358. (doi:10.7717/peerj.17358)

51. Sunagawa S, Desantis TZ, Piceno YM, Brodie EL, Desalvo MK, Voolstra CR, Weil E, Andersen GL, Medina M. 2009 Bacterial diversity and white plague disease-associated community changes in the Caribbean coral Montastraea faveolata. ISME Journal 3, 512– 521. (doi:10.1038/ismej.2008.131)

52. Gignoux-Wolfsohn SA, Vollmer S V. 2015 Identification of candidate coral pathogens on white band disease-infected staghorn coral. PLoS One 10, e0134416. (doi:10.1371/journal.pone.0134416)

53. Voolstra CR et al. 2024 The coral microbiome in sickness, in health and in a changing world. Nat. Rev. Microbiol. 22, 460–475. (doi:10.1038/s41579-024-01015-3)

54. Rojas-Gätjens D, Valverde-Madrigal KS, Rojas-Jimenez K, Pereira R, Avey-Arroyo J, Chavarría M. 2022 Antibiotic-producing Micrococcales govern the microbiome that inhabits the fur of two-and three-toed sloths. Environ. Microbiol. 24, 3148–3163. (doi:10.1111/1462-2920.16082)

55. Ma K-J, Ye Y-L, Li Y-K, Fu G-Y, Wu Y-H, Sun C, Xu X-W. 2025 Polysaccharide metabolic pattern of Cytophagales and Flavobacteriales: A comprehensive genomics approach. Front. Mar. Sci. 12. (doi:10.3389/fmars.2025.1551618)

56. Charpy L, Casareto BE, Langlade MJ, Suzuki Y. 2012 Cyanobacteria in coral reef ecosystems: A review. J. Mar. Biol. 2012, 1–9. (doi:10.1155/2012/259571)

57. Gantar M, Kaczmarsky LT, Stanić D, Miller AW, Richardson LL. 2011 Antibacterial activity of marine and black band disease Cyanobacteria against coral-associated bacteria. Mar. Drugs 9, 2089–2105. (doi:10.3390/md9102089)

58. Sweet M, Burian A, Fifer J, Bulling M, Elliott D, Raymundo L. 2019 Compositional homogeneity in the pathobiome of a new, slow-spreading coral disease. Microbiome 7, 139. (doi:10.1186/s40168-019-0759-6)

59. Suto R, Ishimoto C, Chikyu M, Aihara Y, Matsumoto T, Uenishi H, Yasuda T, Fukumoto Y, Waki M. 2017 Anammox biofilm in activated sludge swine wastewater treatment plants. Chemosphere 167, 300–307. (doi:10.1016/j.chemosphere.2016.09.121)

60. Boeuf D, Eppley JM, Mende DR, Malmstrom RR, Woyke T, DeLong EF. 2021 Metapangenomics reveals depth-dependent shifts in metabolic potential for the ubiquitous marine bacterial SAR324 lineage. Microbiome 9, 172. (doi:10.1186/s40168-021-01119-5)

61. Glasl B, Bourne DG, Frade PR, Thomas T, Schaffelke B, Webster NS. 2019 Microbial indicators of environmental perturbations in coral reef ecosystems. Microbiome 7. (doi:10.1186/s40168-019-0705-7)

62. Webster NS, Reusch TBH. 2017 Microbial contributions to the persistence of coral reefs. ISME J. 11, 2167–2174. (doi:10.1038/ismej.2017.66)

63. Ratcliffe HL, Snyder RL. 1962 Patterns of disease, controlled populations, and experimental design. Circulation 26, 1352–1357. (doi:10.1161/01.CIR.26.6.1352)

64. Stallknecht DE. 2007 Impediments to wildlife disease surveillance, research, and diagnostics. pp. 445–461. (doi:10.1007/978-3-540-70962-6_17)

65. Lewis RE, Davy SK, Gardner SG, Rongo T, Suggett DJ, Nitschke MR. 2022 Colony self-shading facilitates Symbiodiniaceae cohabitation in a South Pacific coral community. Coral Reefs 41, 1433–1447. (doi:10.1007/s00338-022-02292-1)

66. Drake JL, Malik A, Popovits Y, Yosef O, Shemesh E, Stolarski J, Tchernov D, Sher D, Mass T. 2021 Physiological and transcriptomic variability indicative of differences in key functions within a single coral colony. Front. Mar. Sci. 8. (doi:10.3389/fmars.2021.685876)

67. Voolstra CR, Schlotheuber M, Camp EF, Nitschke MR, Szereday S, Bejarano S. 2025 Spatially restricted coral bleaching as an ecological manifestation of within-colony heterogeneity. Commun. Biol. 8, 740. (doi:10.1038/s42003-025-08150-4)

68. Ferdous T, Jiang L, Dinu I, Groizeleau J, Kozyrskyj AL, Greenwood CMT, Arrieta M-C. 2022 The rise to power of the microbiome: power and sample size calculation for microbiome studies. Mucosal Immunol. 15, 1060–1070. (doi:10.1038/s41385-022-00548-1)

69. Silva DP, Epstein HE, Vega Thurber RL. 2023 Best practices for generating and analyzing 16S rRNA amplicon data to track coral microbiome dynamics. Front. Microbiol. 13. (doi:10.3389/fmicb.2022.1007877)

